# Dramatic differences in gut bacterial densities help to explain the relationship between diet and habitat in rainforest ants

**DOI:** 10.1101/114512

**Authors:** Jon G Sanders, Piotr Lukasik, Megan E Frederickson, Jacob A Russell, Ryuichi Koga, Rob Knight, Naomi E Pierce

## Abstract

Abundance is a key parameter in microbial ecology, and important to estimates of potential metabolite flux, impacts of dispersal, and sensitivity of samples to technical biases such as laboratory contamination. However, modern amplicon-based sequencing techniques by themselves typically provide no information about the absolute abundance of microbes. Here, we use fluorescence microscopy and quantitative PCR as independent estimates of microbial abundance to test the hypothesis that microbial symbionts have enabled ants to dominate tropical rainforest canopies by facilitating herbivorous diets, and compare these methods to microbial diversity profiles from 16S rRNA amplicon sequencing. Through a systematic survey of ants from a lowland tropical forest, we show that the density of gut microbiota varies across several orders of magnitude among ant lineages, with median individuals from many genera only marginally above detection limits. Supporting the hypothesis that microbial symbiosis is important to dominance in the canopy, we find that the abundance of gut bacteria is positively correlated with stable isotope proxies of herbivory among canopy-dwelling ants, but not among ground-dwelling ants. Notably, these broad findings are much more evident in the quantitative data than in the 16S rRNA sequencing data. Our results help to resolve a longstanding question in tropical rainforest ecology, and have broad implications for the interpretation of sequence-based surveys of microbial diversity.

## Introduction

When tropical entomologists began systematic surveys of arthropod biomass in rainforest canopies, the dominance of ants in the fauna appeared to be paradoxical. As formulated by Tobin (1991), the problem centered around an apparent inversion of the classic terrestrial ecosystem biomass pyramid: ants were presumed to be predators or scavengers, yet frequently outweighed their putative prey. This biomass “paradox” (Davidson & Patrell-Kim 1996) was partly resolved by evidence from stable isotope analysis that most canopy ants are functionally herbivorous (Cook & Davidson 2006; Douglas 2006; Davidson *et al.* 2003; see also Eilmus & Heil 2009). These ant herbivores feed to a large extent on plant-derived liquid foods, including extrafloral nectar and hemipteran exudates (Davidson *et al.* 2004). But the limited availability of nitrogen in these resources itself poses a dilemma: how do herbivorous canopy ants acquire nitrogen resources that are both abundant and balanced enough in amino acid profile to sustain colony growth?

Insects with nutrient-imbalanced diets frequently rely on bacterial symbioses to complement their nutritional demands (Moran *et al.* 2008; Engel & Moran 2013). Indeed, evidence has been found for specialized associations between bacteria and a number of canopy ant lineages (Russell *et al. In press*)*. Blochmannia* bacteria were among the first described endosymbionts (Blochmann 1888), and appear to play a role in upgrading or recycling nitrogen for their host carpenter ants (genus *Camponotus*), a group frequently found in forest canopies (Feldhaar *et al.* 2007). Specialized extracellular bacteria have long been known to inhabit the morphologically elaborated guts of the new-world arboreal genus *Cephalotes* (Caetano & da Cruz-Landim 1985; Roche & Wheeler 1997; Bution *et al.* 2007). In *Cephalotes,* stability across and correlation with the host phylogeny suggest an important and conserved role for these microbes (Sanders *et al.* 2014), and experimental response to changes in diet suggest that role relates to nutrition (Hu *et al.* 2014). Billen and collaborators showed that several species of the old-world arboreal genus *Tetraponera* have a bacterial pouch at the junction between the mid- and hindgut which houses a dense community of extracellular bacteria (Billen & Buschinger 2000). However, other ant lineages that have been surveyed, including the invasive fire ant, show less evidence for specialized bacterial associations (Lee *et al.* 2008; Ishak *et al.* 2011; Sanders *et al.* 2014).

In the first major comparativ analysis that systematically surveyed bacteria across ants, including representatives of two-thirds of known ant genera, Russell and colleagues detected a systematic relationship between herbivory (defined by stable isotope composition) and presence of an ant-specific lineage of alphaproteobacteria related to the genus *Bartonella* (Russell *et al.* 2009). As previously reported in *Tetraponera* (Stoll *et al.* 2007), Russell *et al.* were able to amplify the *nifH* gene from some of these ant specimens, though no nitrogenase activity has yet been reported in functional assays for nitrogen fixation. But even without direct evidence of a functional relationship, the distribution of ant-specific bacteria across lineages likely to have independently-evolved herbivorous diets makes a compelling case for a generalized role for bacteria in facilitating herbivory in ants.

Much of the research that has been done to describe insect-associated bacterial communities, especially since the advent of high-throughput next-generation sequencing, suffers from a common limitation: a lack of context as to the absolute abundance of the microbes being surveyed. PCR-based microbial community profiling techniques, including cloning and Sanger sequencing, restriction fragment polymorphism analysis, and next-generation amplicon sequencing, almost always start with an amplification step to produce many copies of the original template DNA. The resulting libraries retain almost no information about starting template abundance, and even information about the relative abundance among different taxa is subject to biases (Engelbrektson *et al.* 2010). The amplification step can be subject to contamination, especially for samples (like ants) with very low starting DNA concentrations (Salter *et al.* 2014; Lukasik *et al.* 2016; Hu *et al. In Press*; Russell *et al. In Press*). Even in the absence of contamination, the biological implications of very low density bacterial communities are likely to be substantially different than for symbionts, like the nutritional endosymbionts of *Camponotus* ants (Wolschin *et al.* 2004), that are present in their hosts at very high numbers. Without additional information about absolute abundance, it can be difficult to draw meaningful biological conclusions from diversity.

Insects known to rely on bacterial symbionts for nutrient complementation also tend to support relatively high densities of those symbionts (Schmitt-Wagner *et al.* 2003; Martinson *et al.* 2012; Engel & Moran 2013). Here, we examined how bacterial abundance varies across a wide range of ant species in a tropical rainforest, and whether ants at lower trophic levels support more bacteria. We used two independent methods, quantitative PCR and fluorescence microscopy, to assess absolute abundance of cells while also gaining insight into their localization and morphology. We then contrasted these results to the patterns that could be detected in amplicon sequencing data alone. Our findings reveal surprising diversity in the nature and density of these associations, providing critical context for understanding the roles that microbes may play in these important members of the rainforest ecosystem.

## Methods

### Field collections

We performed primary collections in July-August 2011 at the Centro de Investigación y Capacitación Rio Los Amigos (CICRA) in southeastern Peru, approximately 80 km west of Puerto Maldonado. CICRA contains a mix of primary and secondary lowland tropical forest. We collected opportunistically from most available habitat types at the station, finding ants primarily visually but also using a mix of baits to recruit workers in some cases. To ensure that individuals came from the same colony, we took workers from within nests when possible; but when nests were inaccessible or could not be found, multiple workers were taken from the same foraging trails. In all cases, we brought live workers and/or nest fragments back to the field station for processing. Each colony was processed within 24 hours of collection.

When numbers allowed, we preserved tissues for nucleic acid analysis, stable isotope analysis, FISH microscopy, and morphology. First, workers were sacrificed by brief (1-5 minutes) immersion in 97% ethanol. They were subsequently surface sterilized in 0.5% sodium hypochlorite solution for approximately 1 minute, then rinsed twice in sterile PBS buffer. For preservation of nucleic acids, the midgut and hindgut of worker ants were dissected with sterile forceps in clean PBS buffer and preserved in RNAlater, one ant per vial. One gastrointestinal tract per colony was also completely dissected and visualized immediately using fluorescence microscopy (see below). The heads, legs, and mesosomas from these dissected individuals were preserved together in 95% ethanol in a separate tube for analysis of stable isotopes.

To preserve for subsequent fluorescence microscopy, we semi-dissected whole worker gasters to expose internal tissues. These were fixed in 4% PBS-buffered paraformaldehyde and preserved in molecular grade ethanol, as described in detail in Supplemental Methods.

Any remaining workers were preserved whole in 95% ethanol for morphological identification.

In addition to the primary collection at CICRA, we collected a secondary set of specimens in August 2013 for additional FISH microscopy. These collections took place in secondary forest at the Villa Carmen field station near the town of Pilcopata, Cusco province, Peru, approximately 140 km west of the primary collection site.

Ants were morphologically identified by Dr. Stefan Cover of the Museum of Comparative Zoology at Harvard University.

### Microscopy

For most colonies, we visualized a single dissected worker gut in the field using SYBR Green fluorescence microscopy. Guts dissected as above were placed on a glass slide, covered with a 1:100 mixture of SYBR Green and VectaShield mounting medium, and torn open using forceps to expose the contents of the midgut and ileum. Slides were covered with a glass coverslip and sealed with clear nail polish, then visualized on an AmScope epifluorescence microscope (model number FM320TA) powered by a portable generator. Putative bacterial cells were identified by size and morphology, and the abundance estimated using a roughly logarithmic visual scale (0 = no visible bacterial cells, 1 = tens, 2 = hundreds, 3 = thousands, 4 = tens of thousands; see Fig. S1). Representative photomicrographs for each colony were taken with a digital camera.

Preserved ant tissues proved especially difficult to use for FISH microscopy relative to tissues from other insects, rapidly losing morphological structure when fixed in only acetone or ethanol, and displaying high levels of autofluorescence. For convenience, we have provided detailed protocols as supplemental material. Briefly, fixed semi-dissected ant gasters were rehydrated in PBSTx solution, then bleached with 80% ethanol – 6% hydrogen peroxide solution for several days in order to decrease tissue autofluorescence (Koga *et al.* 2009) and then rehydrated again and carefully dissected when necessary. For whole-mount FISH, specimens were rehydrated in buffer, dissected further if necessary, washed in hybridization solution, and hybridized with a solution containing FISH probes and DAPI. Hybridized samples were then washed and mounted in an antifade medium on a slide for visualization. Specimens for tissue sections were dehydrated in acetone before embedding in glycol methacrylate resin (Technovit), and 1-2 *μ*m sections cut on a microtome. These sections were hybridized with a solution containing FISH probes and DAPI, then washed and visualized under an antifade medium.

### Nucleic acids analysis and quantitative PCR

We extracted DNA from individual dissected and RNAlater-preserved guts using the PowerSoil 96-well DNA extraction kit from MoBio, using individual extractions from three worker guts per colony when possible. First, we added 1 volume of sterile molecular-grade water to tubes containing dissected guts to help redissolve any precipitated ammonium sulfate. Tubes were vortexed several times at room temperature until any visible precipitate had dissolved, then spun in a microcentrifuge at 10,000 xg for 10 minutes to pellet cells and tissues. We removed the supernatant and replaced it with 200*μ*L buffer C1 from the PowerSoil extraction kit, vortexed at maximum speed for 15 seconds to resuspend tissues, and transferred this solution to the extraction plates. From there, we proceeded with the extraction according to the manufacturer’s protocol.

We quantified total extracted DNA using PicoGreen dsDNA quantification reagent (Thermo Scientific), following the manufacturer’s protocol for 384-well microplate formats (Thermo Scientific, 2007). Due to limited quantities of eluted DNA, the protocol was modified slightly: rather than mixing equal volumes sample and PicoGreen reagent solution, 10 *μ*L of sample solution was added to 30 *μ*L quantitation reagent, diluted correspondingly with molecular grade water. Each sample was measured in triplicate on the same 384-well microplate. Plates were read on a Spectramax Gemini XS fluorescence plate reader, and standard curves fit in SOFTmax PRO (Molecular Devices, Inc.). The mean of the three replicates was taken as the DNA concentration for each extraction.

PCR quantitation of bacterial 16S rRNA genes copies was performed with SYBR Green chemistry (PerfeCTa SYBR Green SuperMix, Quanta Biosciences) using the primers 515F and 806R (Caporaso *et al.* 2011), each at 250 pM. This primer pair was chosen to permit direct comparison of qPCR values with Illumina-sequenced amplicons of the same locus. Two microliters of extracted DNA were used per 20 *μ*L amplification reaction in 96-well plates. Reactions were performed on a Stratagene MX3000p realtime thermocycler, using 40 iterations of the following three-step cycle: 45 seconds denaturation at 94° C, 60 seconds annealing at 50°, and 90 seconds extension at 72°. In addition, a 3 minute initial denaturation at 94° and a post-amplification denaturation curve were performed. To increase measurement accuracy, each sample was run at least twice, with each replicate occurring on a separate PCR plate. These technical replicates of each individual were averaged before further analysis, and figures illustrating these data present the median value among individuals in a colony. For absolute quantification, we included in triplicate a 1:10 serial dilution standard curve generated from linearized plasmids containing a full-length E. coli 16S rRNA gene. Due to background amplification from 16S rRNA genes present in reagents, some amplification was observed at high cycle numbers in no-template controls (mean NTC amplification estimated at 85 copies / *μ*L). The mean background amplification from three no-template controls per plate was subtracted from each sample on that plate, and samples below this limit of detection normalized to 1 copy per *μ*L. To test for specificity of qPCR primers, we also prepared a standard curve of eukaryotic 18S rRNA genes amplified from an ant (*Cephalotes varians*).

### 16S rRNA amplicon sequencing

To enable comparisons between abundance-based and diversity-based 16S analyses, aliquots of the DNA used for molecular quantitation were also sequenced using standard bacterial 16S rRNA gene amplicon sequencing protocols as part of the Earth Microbiome Project (http://www.earthmicrobiome.org). Briefly, the V4 region of the 16S rRNA gene was PCR-amplified in triplicate using the primers 515fB (GTGYCAGCMGCCGCGGTAA) and 806rB (GGACT ACNVGGGTWT CT AAT) (Caporaso *et al.* 2012; Walters *et al.* 2016). Pooled amplicons were then sequenced on an Illumina MiSeq instrument at the Center for Microbiome Innovation at the University of California, San Diego.

Per standard EMP processing protocols, sequences were uploaded to Qiita (https://qiita.ucsd.edu) for quality filtering and demultiplexing, and the demultiplexed forward reads downloaded for further analysis in QIIME v.1.8.1 (Caporaso *et al.* 2010b). Full analysis scripts are provided in supplemental information; but briefly, reads were chimera-checked and clustered into 97% OTUs using the vsearch (Rognes *et al.* 2016) implementation of the UPARSE pipeline (Edgar 2013). Taxonomy was assigned to OTUs using uclust against the Greengenes 97% OTU database, representative sequences aligned using PyNAST (Caporaso *et al.* 2010a), a phylogenetic tree estimated with FastTree (Price *et al.* 2009), and beta-diversity distances calculated using unweighted UniFrac (Lozupone & Knight 2005).

### Isotopic analysis

To estimate the relative trophic position of the ant colonies in this study, we analyzed ethanol-preserved tissues using stable isotope ratio mass spectrometry. Heads and mesosomas from the individuals used for gut dissections (three or more individuals per colony) were preserved in a separate vial of 95% ethanol to minimize the isotopic contribution of materials from the gut. For each colony analyzed, these tissues were dried overnight at 60° C, ground into powder with a mortar and pestle, and ~ 5 mg of powder placed in a silver foil capsule. These were combusted and analyzed for *∂*15N at the Boston University Stable Isotope Laboratory.

### Statistical analysis

We tested the correlation of both visual and quantitative PCR estimates of bacterial abundance using linear and generalized linear mixed models with the lme4 package in R. Because visual estimates corresponded roughly to a step function with respect to qPCR estimates (Fig. S2), we treated these data as presence / absence, with visual estimates of 0 or 1 corresponding to ‘absent’ and 2 - 4 corresponding to ‘present’. Bacterial presence per colony was modeled using logit-linked binomial regression with the fixed effects of ∂15N, habitat, and DNA concentration (as a proxy for total host and microbe biomass), treating host genus as a random effect. Quantitative PCR estimates of (non DNA-concentration normalized) 16S rRNA gene abundance per individual were modeled with a linear mixed model using the same fixed effects, but using colony nested within host genus as random effects. For both GLMM and LMMs, Akaike Information Criteria and likelihood ratio tests were used to select the best model.

## Results

### Collections

At CICRA, we collected data for a total of 97 colonies from 29 genera. Of these, 54 were collected from arboreal and 38 from terrestrial habitats. Voucher specimens for each colony have been deposited with the Centre de Ecología y Biodiversidad (CEBIO) in Lima, Peru and the Museum of Comparative Zoology (MCZ) in Cambridge, MA, USA. Detailed collections information can be found in Table S1 and in the metadata associated with these samples in the Earth Microbiome Project (https://qiita.ucsd.edu/study/description/10343).

### Visual microscopy survey

Most ant guts surveyed by SYBR Green fluorescence microscopy did not harbor identifiable bacterial cells (N = 59; Fig. 1a). In these guts, although host nuclei were clearly stained and highly fluorescent (Fig. S3a), and gut contents could be seen spilling from the punctured gut under light microscopy and occasionally via autofluorescence (Fig. S3b), there were no visible DNA-containing cellular structures in the size range typical of bacteria. Several of the dissected guts did contain just a few apparent bacterial cells (visual rubric score of 1, N = 8). All individuals examined from the abundant and typically ground-nesting genera *Solenopsis* and *Pheidole* fell into these categories, as did all of the leaf-cutting ants, most individuals from ground-dwelling genera formerly grouped in the subfamily Ponerinae (including *Ectatomma* and *Pachycondyla*), and most of the individuals from the arboreal genera *Azteca, Crematogaster,* and *Pseudomyrmex.* Individuals from genera that typically hosted few or no apparent bacterial cells (such as *Azteca* and *Crematogaster*) did occasionally contain high densities, although the reason for this variability is unclear.

**Fig 1:**
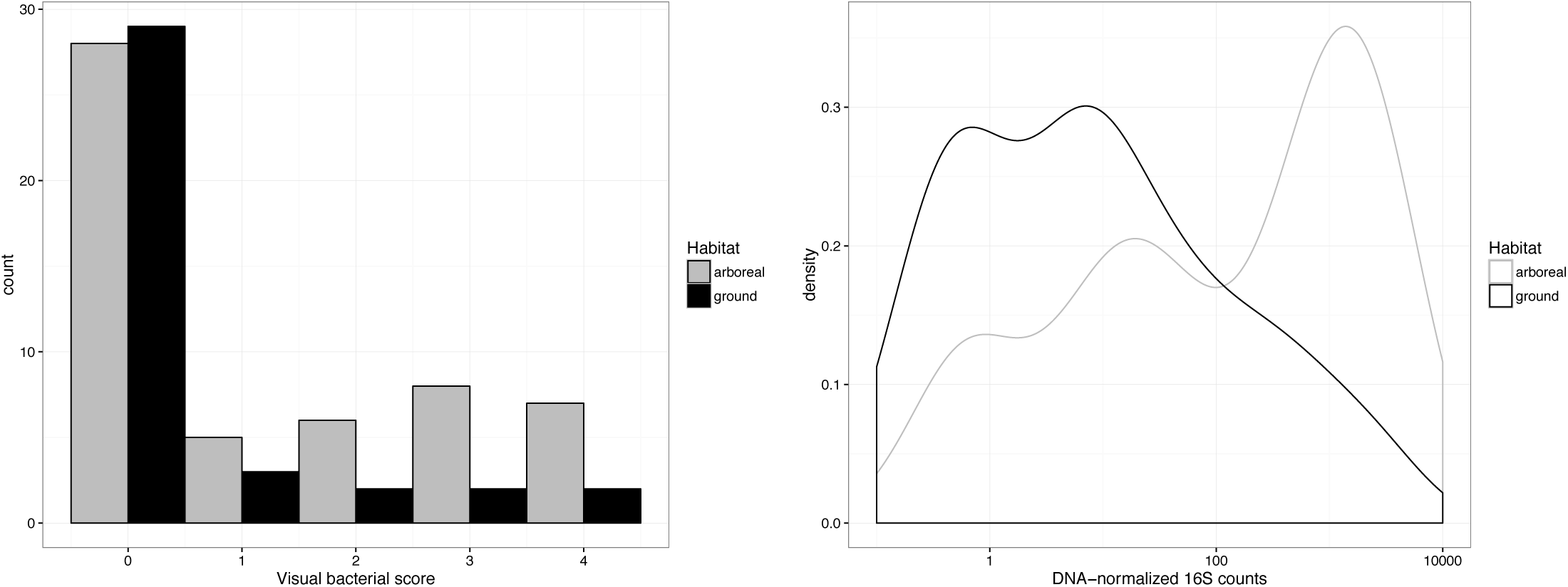
a) Histogram of visual bacterial abundance estimates from *in situ* fluorescence microscopy. Estimates followed a roughly logarithmic scale (0 = no visible bacterial cells, 1 = tens, 2 = hundreds, 3 = thousands, 4 = tens of thousands; see Fig. Sn1). b) Kernel density plot of normalized bacterial abundance estimates from quantitative PCR.

By contrast, the density of bacterial cells in other ant guts was striking. Cell densities in guts of the abundant arboreal taxa *Camponotus*, *Cephalotes*, and *Dolichoderus,* were often so high that out-of-plane fluorescence inhibited photography using our field microscopy equipment (Fig. S3c). Moderate to high bacterial cell densities were frequently observed in army ants, including the ecitonine genera *Labidus* and *Eciton* as well as the cerapachyne genus *Acanthostichus* (visual scores 2-4, N = 4), although these genera also frequently appeared devoid of bacteria (visual scores 0-1, N = 4). The single individuals examined of myrmicine genera *Basiceros* and *Daceton* both hosted fairly high densities of putative bacterial cells.

At least three genera appeared to harbor bacterial cells in bacteriocytes localized to the midgut. These specialized cells were very clearly visible in the individuals we examined from *Camponotus,* appearing in the SYBR Green gut squash preps as swaths of bright green patches intercalated with midgut cells (Fig. S3e,f). The process of puncturing the gut always disrupted a number of these bacteriocytes, spilling large numbers of the intracellular bacteria into the surrounding mounting medium. These cells were morphologically distinct, often quite large, and sometimes showed very long cells with intracellular DNA aggregation under high magnification suggestive of polyploidy (Fig. S3d; Fig. 2).

**Fig. 2:**
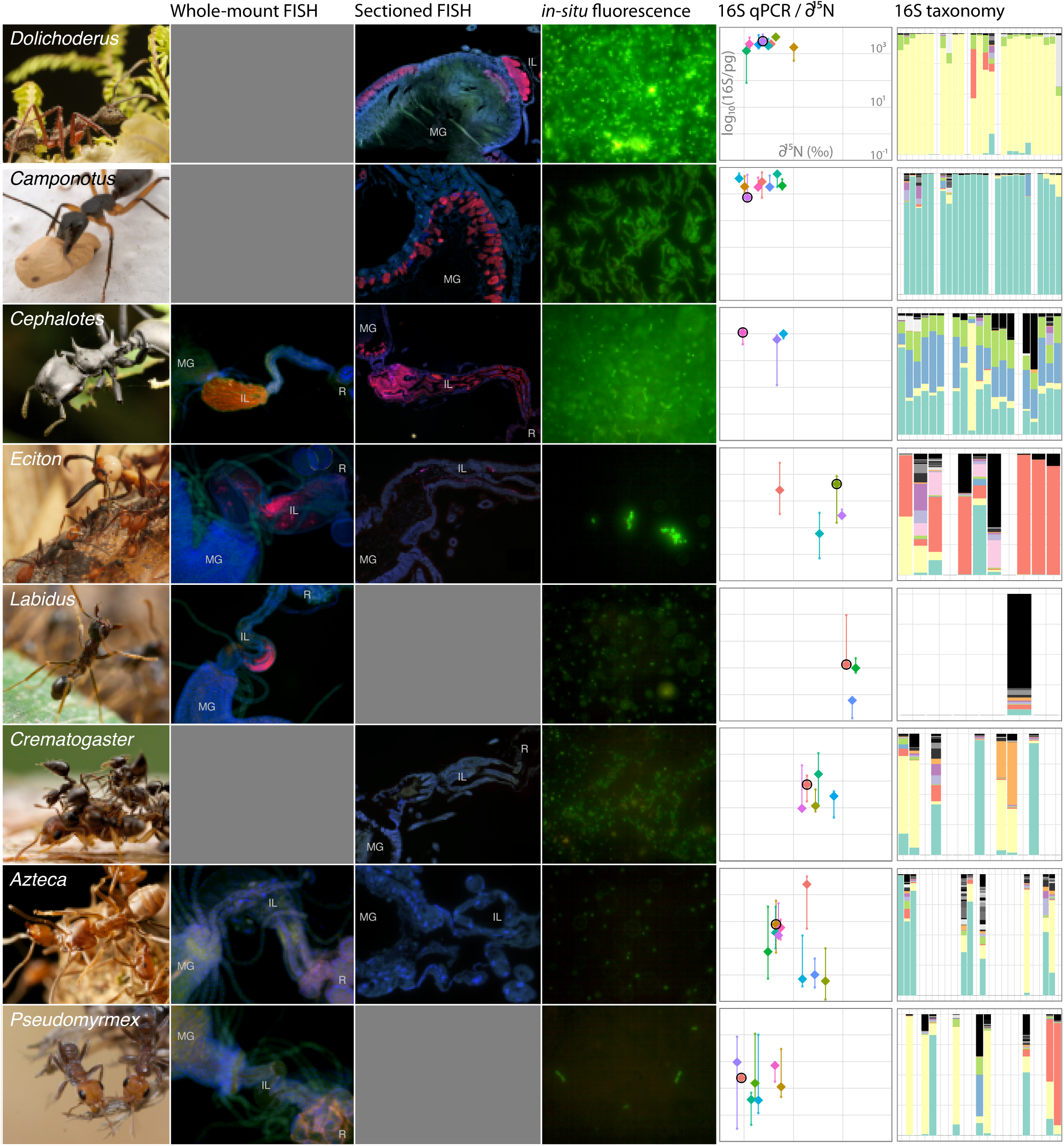
Summarized microscopic and molecular evidence of gut bacterial abundance in eight common Peruvian ant genera. Column 1: Photographs and genus names (© JGS). Column 2: False-color FISH micrograph of whole-mount dissected guts. Tissue autofluorescence is visible in all three channels (blue, green, and red). DNA is stained with DAPI in the blue channel, and the universal bacterial probe Eub338 is hybridized in the red channel. MG: midgut; IL: Ileum; R: rectum. (© PL). Column 3: False-color FISH micrographs of resin-embedded tissue sections. Note bacteria present both putatively intracellularly (midgut wall) and extracellularly (in lumen of ileum) in *Dolichoderus,* and only intracellularly in *Camponotus.* Colors and labels as in Column 2 (© PL). Column 4: SYBR Green fluorescence micrographs of bacteria from gut squashes. All images are uncropped and taken under identical magnification (40X objective). *Dolichoderus* image taken of bacteria from hindgut lumen (© JGS). Column 5: normalized log_10_ bacterial 16S rRNA gene copy number by ∂^15^N isotope ratio. Large diamonds represent median values per colony, with small points representing individuals. Lines indicate range of values observed for each colony. Each graph is on the same scale. The colony from which the SYBR Green micrograph from Column 4 was taken is indicated by a black circle. Column 6: Class-level taxonomic composition of individual samples, per host genus. Samples that were included on the sequencing attempt but that did not yield successful sequencing libraries are represented by blank columns. Ten most abundant classes are colored as in Fig. S9; others randomly assigned grey values.

We also observed morphologically similar host cells, or putative bacteriocytes, in the midguts of individuals from one of the *Myrmelachista* specimens we examined (Fig. S3g,h). As in *Camponotus,* these appeared as fairly distinct bright green patches distributed around the midgut. The putative bacterial cells in these individuals presented as relatively large rods, though without the obviously anomalous morphologies frequently observed in *Camponotus* bacteria.

Individuals of many *Dolichoderus* also exhibited patterns of DNA fluorescence staining consistent with intracellular bacteria localized to the midgut. Unlike in *Camponotus,* where bacteriocytes were visible as clearly bounded cells, *Dolichoderus* midguts stained with SYBR Green appeared shrouded in a uniform green glow, largely obscuring the distinct host nuclei typically visible in other midguts. At higher magnification, these could be resolved as masses of morphologically unusual cells. Like the *Blochmannia* bacteria we observed erupting from *Camponotus* bacteriocytes, the putative intracellular bacteria in *Dolichoderus* were relatively large (Fig. 2). They were also often branched, again consistent with deficiencies in cell division and cell wall synthesis observed in other intracellular bacteria of insects (McCutcheon & Moran 2011). Along with these highly unusual cells, some *Dolichoderus* specimens exhibited high densities of smaller, coccoid bacterial cells. In at least one specimen for which we separately dissected midgut and hindgut compartments, these coccoid cells appeared localized to the hindgut (data not shown).

Targeted microcopy using fluorescently labeled universal bacterial 16S rRNA probes (FISH microscopy) supported our inferences from field-based SYBR Green microscopy. Whole-mount and resin sectioned guts from *Azteca, Pseudomyrmex,* and *Crematogaster* showed no evidence of bacteria, while several specimens from the army ants *Labidus* and *Eciton* showed small populations of bacterial cells localized to the hindgut epithelia (Fig. 2). By contrast, very large populations of bacteria could be readily seen in sections from *Camponotus, Cephalotes,* and *Dolichoderus.* In specimens of *Camponotus japonicus* (not collected from Peru, but used as a representative sample from this well-studied genus), *Blochmannia* bacteria can be clearly observed in bacteriocytes interspersed among the midgut epithelia. In *Dolichoderus,* bacteria are visible forming a relatively uniform layer among midgut cells close to where they border the hemolymph, as well as forming a dense mass in the pylorus and upper part of the ileum. In *Cephalotes,* bacterial cells form an aggregate that almost entirely fills the enlarged and highly folded ileum, as has been described previously through visible light and electron microscopy (Bution *et al.* 2007). Uniquely in *Cephalotes,* we also observed fluorescence indicating masses of bacterial cells in the distal part of the midgut lumen.

It should be noted that, for all ant specimens examined, high levels of tissue autofluorescence interfered with the relatively weak signal from monolabeled FISH probes. Despite the hydrogen peroxide pre-treatment which was shown to effectively reduce autofluorescence in other insects (Koga *et al.* 2009), in ant tissues autofluorescence was observed over a wide range of wavelengths, and was especially pronounced in the blue to green range. Tissues lined with chitin (such as the crop and the rectum) displayed comparatively elevated levels of autofluorescence at longer wavelengths. Fat bodies and malpighian tubules also showed especially strong autofluorescence. Consequently, special care should be taken when interpreting FISH hybridizations from ant guts.

### Quantitative PCR

Estimation of bacterial abundance via quantitative PCR targeting 16S rRNA genes largely corroborated visual estimates from field-based SYBR Green microscopy, with arboreal taxa tending to have higher counts (Fig. 1b). The per-colony median 16S rRNA gene concentration correlated well with visual abundance estimates (Spearman’s ρ = 0.44, *p* << 0.001), especially after normalizing by DNA concentration, a proxy for quantity of extracted tissue (Spearman’s ρ = 0.53, *p* << 0.001). Colonies with a visual abundance score of 0 or 1 had DNA-normalized 16S rRNA gene concentrations statistically indistinguishable from one another, but significantly lower than colonies with visual scores of 2-4 (Fig. S2).

Estimates of 16S copy number correlated strongly with DNA concentration (Fig. S4a). For many smaller-bodied ant species, this meant that 16S rRNA gene quantities were below the detection threshold set by background amplification, or about 85 copies per *μ*L. The lower bounds of detection may also have been affected by amplification of host 18S rRNA gene molecules, which our primer set amplified with much lower affinity than bacterial 16S rRNA genes (18S rRNA gene standard curve concentrations were underestimated by a factor of between 10^4^ and 10^5^ with the 16S rRNA gene primers used). Thus, relative differences between high-and low-abundance samples are likely to be underestimated by this method.

Despite these limitations, qPCR estimates revealed dramatic differences in the median bacterial abundance in colonies of different ant genera (Fig. 3; Fig. S5). To convey a sense of relative bacterial abundance independent of host body size, we use [DNA]-normalized values, expressed in 16S rRNA gene copies per pg DNA (Fig. S4b). This normalization will tend to underestimate relative bacterial abundance in samples for which bacterial DNA makes up a very large proportion of total DNA as it asymptotically approaches the ratio of 16S rRNA gene copies per bacterial genome, but has the advantage of being insensitive to extraction efficiency. These normalized abundances ranged from a minimum of 0.081 copies/pg in one ponerine colony, in which the median individual concentration was below the limit of detection despite a relatively high DNA concentration, to a maximum of 5537 copies/pg in a colony of *Camponotus.* The extremes were not dramatic outliers: the first and third quartiles were separated by more than two orders of magnitude (1Q: 3.77, 3Q: 931 copies / pg). Consistent with field microscopy observations, most genera we sampled had very low normalized bacterial abundances: only 10 of the 29 had median 16S rRNA gene counts above 100 copies / pg.

**Fig. 3:**
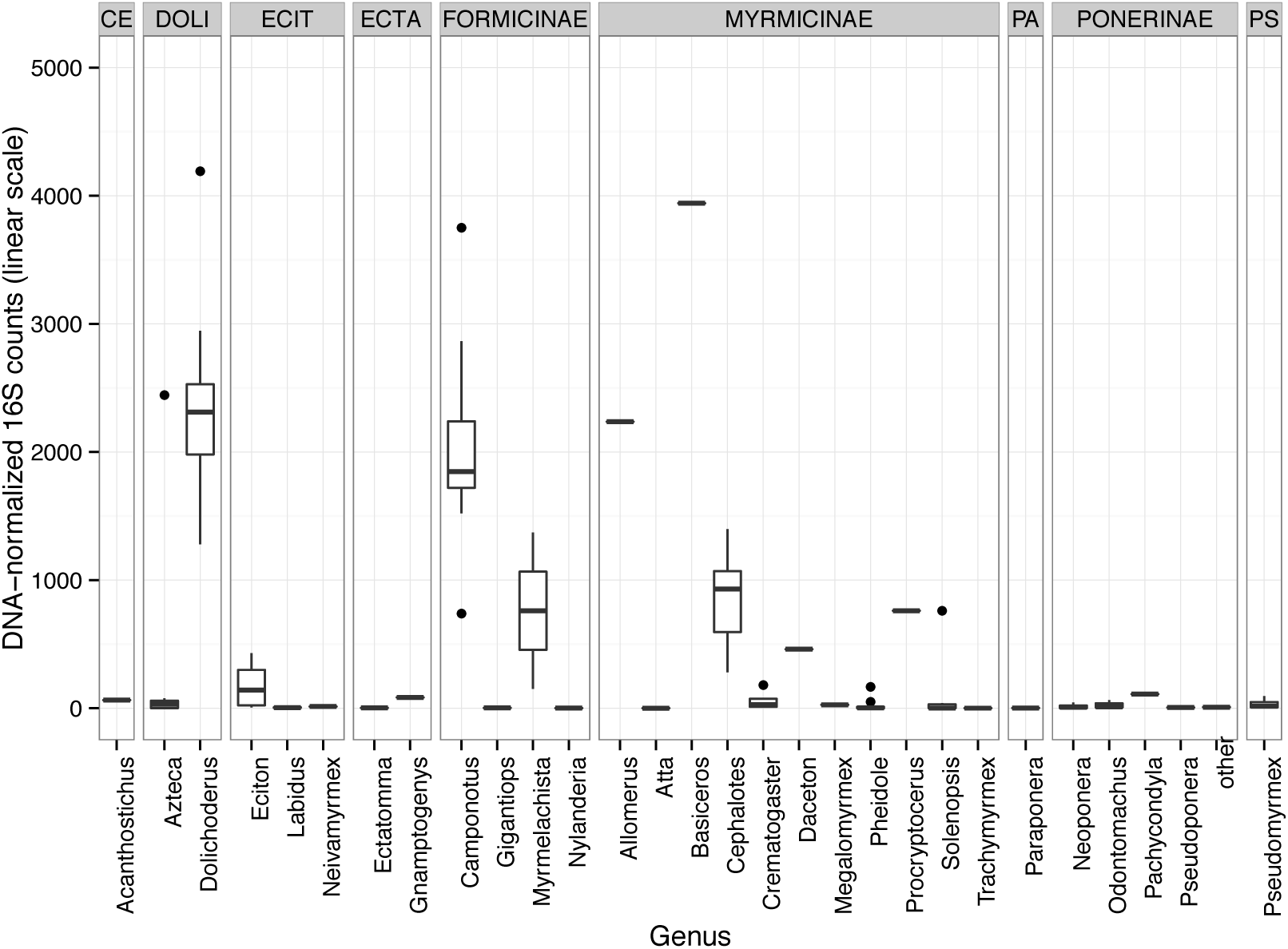
Normalized bacterial abundances by genus. Data shown are 16S rRNA qPCR counts, minus mean non-template control counts, divided by total DNA concentration. Each data point represents a single colony, taken as the median of three individuals. CE = Cerapachyinae; DOLI = Dolichoderinae; ECIT = Ecitoninae; ECTA = Ectatomminae; PA = Paraponerinae; PS = Pseudomyrmicinae.

Genera with high bacterial normalized abundances tended to host consistent numbers among colonies. Colonies of the arboreal genera *Cephalotes, Camponotus,* and *Dolichoderus* all had consistently high normalized median 16S rRNA gene abundances, with maximum and minimum values within an order of magnitude, despite relatively large numbers of colonies sampled (Fig. S6). *Camponotus* and *Dolichoderus* colonies had somewhat higher normalized median 16S abundances than did *Cephalotes* (medians per genera of 2031 and 2257 copies / pg vs 931.0 copies / pg, respectively). Of the twelve other genera for which we had sampled at least two colonies, in ten the average normalized bacterial abundances varied between colonies by more than an order of magnitude. One of the remaining genera, *Myrmelachista,* hosted relatively high numbers of bacteria (151 and 1371 16S rRNA copies / pg in the two examined colonies; this was also noted using field microscopy), while both *Megalomyrmex* colonies we examined had fairly low numbers (26 and 27 16S rRNA copies / pg)

### 16S amplicon sequencing diversity analysis

Of 288 dissected ant gut samples, 169 yielded more than the 10,000 sequences we chose as a rarefaction cutoff. Generally, and has been reported previously (Rubin *et al.* 2014), these samples derived from samples with higher absolute counts of the 16S rRNA gene as measured by qPCR (Fig. S7). Beta-diversity analyses using the unweighted UniFrac metric did not show clear patterns of microbial community turnover with respect to ∂^15^N ratio or host ant arboreality (Fig. 4), but did show some grouping by host taxonomy (Fig. S8).

**Fig. 4:**
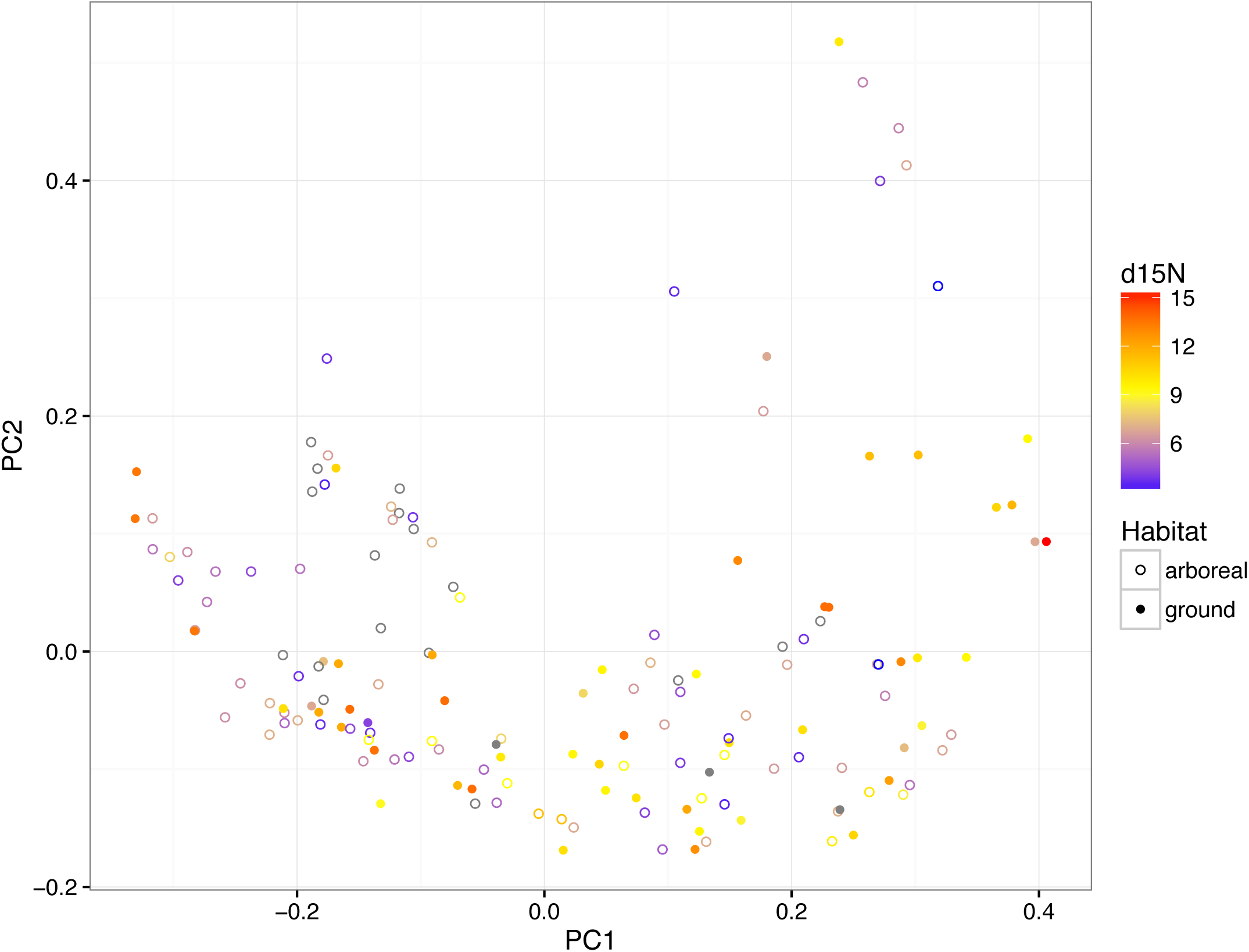
Principle Coordinates analysis of unweighted UniFrac distances among samples. Individual samples are colored by stable nitrogen isotope ratio with shape indicating habitat, filled circles corresponding to ground-dwelling.

Microbial taxonomic profiles inferred by 16S rRNA sequencing were more consistent across individuals in host genera with higher bacterial abundance (Fig. 2, Fig. S9). Microbial diversity within colonies also correlated with estimates of bacterial abundance. Colonies with high median qPCR estimates of bacterial abundance had lower median pairwise UniFrac distances (Fig. S10). The strength of this correlation was higher for [DNA]-normalized 16S counts than for raw 16S counts (*r*^2^ = 0.281 for log 10 normalized raw 16S counts, *r*^2^ = 0.373 for log_10_ [DNA]-normalized 16S rRNA counts), as smaller ants with large quantities of bacteria for their size also hosted consistent communities.

Sequence data are available for analysis on the Earth Microbiome Project portal on Qiita, study number 10343 (https://qiita.ucsd.edu/study/description/10343), and deposited in the EMBL-EBI European Nucleotide Archive, accession number ERP014516.

### Correlation of bacterial abundance with ecological variables

To determine whether gut bacterial abundance correlates significantly with host ecology, we fit linear mixed models to visual and qPCR estimates of abundance, using host habitat (arboreal or terrestrial) and relative trophic position (inferred by ∂^15^N ratio, Fig. S11) as fixed effects and genus and colony as random effects. As has been previously described from a similar sample of ants at a nearby site (Davidson *et al.* 2003), arboreal ants showed signatures of feeding at significantly lower trophic levels; ∂^15^N ratios differed significantly by habitat (two-tailed students t test *p* << 0.001), with a mean of 6.57‰ in arboreal and 10.5‰ in terrestrial ants.

Fitting a generalized linear mixed model to presence or absence of bacteria in our field microscopy survey indicated that bacterial presence was significantly associated with habitat (*p* = 0.0187) as well as trophic position (*p* = 0.0451), but that the directionality of association with trophic position was opposite in arboreal and terrestrial habitats (interaction *p =* 0.0155). In arboreal ants, herbivorous colonies – those with lower ∂^15^N ratios – were more likely to host visible bacterial cells. But among terrestrial ants, the opposite was the case: bacteria tended to be found in more carnivorous ants (Fig. S12a). The model with this interaction term had a significantly better fit and lower Akaike Information Criterion values than models without it (Table S2).

Quantitative estimates of bacterial abundance via qPCR gave similar results. We fit linear mixed models of absolute bacterial 16S rRNA gene quantity with DNA concentration, habitat, and relative trophic position as fixed effects, taking colony nested within genus as random effects. As expected, 16S rRNA gene quantity correlated strongly with DNA concentration (*p* < 0.0001). Consistent with our findings above, bacterial abundances were higher in arboreal ants than in terrestrial ants (*p* = 0.0043) and correlated with relative trophic position (*p* = 0.0061), but the direction of correlation between bacterial abundance and relative trophic position differed in each habitat (*p* = 0.0238; Fig. S12b). As with the microscopy data, the model with an interaction term had significantly better fit and lower AIC (Table S3). To ensure that these results were not due to differences in collecting methodology, we also evaluated models including collection method as an additional fixed effect; models including this variable were not significantly better and had higher AIC than models without (Table S4). In the best model that included collection method as an effect (lmm.3 in Table S4), samples collected from baits did not have significantly different 16S rRNA gene quantities than those collected foraging (*p* = 0.5911) or from nests (*p* = 0.6313).

## Discussion

Our findings support the hypothesis that symbioses with bacteria are systematically important to the dominance of ants in the tropical forest canopy (Davidson *et al.* 2003; Cook & Davidson 2006; Russell *et al.* 2009), reflected by variation in normalized bacterial abundance across several orders of magnitude. Using two independent methods of characterizing bacterial abundance, we found bacteria to be both more abundant in arboreal ants, and a predictor of herbivory among arboreal ants. Surprisingly, most of the ants we surveyed had very few bacteria. Estimates of bacterial abundance were also much more tightly associated with these ecological variables than were estimates of bacterial beta diversity as measured with standard 16S rRNA sequencing protocols, highlighting the benefit of microbial quantitation in broad surveys of microbiota. Together, our findings present the beginnings of a systematic framework for understanding the relationship between diet and bacterial symbiosis in ants.

### Bacterial abundance and ant ecology

Our findings support a relationship between bacteria and herbivory in canopy ants: almost all of the ants with very high normalized bacterial abundances were canopy ants at the herbivorous extreme of the ∂^15^N scale, and the correlation between ∂^15^N isotope ratios and bacteria was significant for both microscopic and molecular measures of bacterial abundance. But while the ants with the highest numbers of bacteria appeared to be mostly herbivorous, maintaining such high titers in worker guts does not seem to be essential to ant life in the canopy, or even to highly specialized herbivory. In our visual survey of ant guts, the distribution of bacterial abundance was strongly bimodal, with many arboreal individuals we surveyed not obviously hosting any bacterial cells at all (Fig. 1). Our qPCR-based estimates of normalized bacterial abundance in arboreal ants were similarly bimodal. Ants represented by the lower peak of this distribution appear to be utilizing fundamentally different approaches to the challenge of acquiring nitrogen in the canopy.

The high-abundance peak of bacterial distribution was composed almost entirely of ants belonging to one of three taxa – *Camponotus, Cephalotes* (and its sister genus *Procryptocerus),* or *Dolichoderus* – that have previously been linked with bacterial symbioses. Of these, *Camponotus* symbioses are the best studied, with gamma-proteobacterial *Blochmannia* endosymbionts implicated in the recycling/upgrading of nitrogen from urea into essential amino acids (Feldhaar *et al.* 2007). As expected, *Blochmannia* gamma-proteobacteria dominated the 16S rRNA sequence profiles of the *Camponotus* in our study (Fig 2). The experimental evidence for a nutritional role in *Cephalotes* symbionts is to this point more limited (Jaffe *et al.* 2001). They host a moderately complex bacterial community in their gut lumen, comprising at least one species of Verrucomicrobia and several species of alpha-, beta-, and gamma-proteobacteria. The *Cephalotes* gut community is both consistent and phylogenetically correlated across the genus (Sanders *et al.* 2014), a pattern we also recovered in our 16S rRNA sequencing for this study (Fig 2), and has shown some sensitivity to changes in diet (Hu *et al.* 2014). Little is known about the bacterial associates of *Dolichoderus* beyond a handful of 16S rRNA gene clones sequenced from a few individuals as part of other studies (Stoll *et al.* 2007; Russell *et al.* 2009; Anderson *et al.* 2012), but the sequence similarity of these clones to others sequenced from herbivorous ants has led some to speculate that they play a similar functional role. Here, we found that virtually every *Dolichoderus* individual in our study was dominated by sequences classified as belonging to the alpha-proteobacterial class Rhizobiales, consistent with previous studies; as well as lower but consistent numbers of sequences classified as the beta-proteobacterial class Burkholderiales – suggesting that, at least among the arboreal Neotropical *Dolichoderus* we sampled, bacterial communities are highly conserved by identity as well as by quantity. Together, these three genera, all well-represented in our collection, are responsible for virtually all of the correlation we observed between bacteria and herbivory: excluding them, there was no significant relationship between ∂^15^N isotope ratio and herbivory.

We posit that high bacterial abundances in these genera are necessary to sustain large nutrient fluxes. Despite major differences in the identity and physiology of their symbiotic associations, they have converged on a similar density. We measured median DNA-normalized 16S rRNA gene abundances in these genera that were almost all within an order of magnitude of one another. Abundances within *Cephalotes* were somewhat lower than in *Dolichoderus* and *Cephalotes,* though polyploidy in endosymbiotic bacteria and differences in per-genome 16S rRNA gene copy number make accurate extrapolation to absolute cell counts uncertain. What is certain is that workers from these three genera consistently maintain bacterial densities that are orders of magnitude greater than those found in most other ants. That such consistent associations have arisen independently in these three lineages, each with markedly herbivorous stable isotope signatures, lends additional credence to the hypothesis that bacteria play an important and convergent functional role in these canopy ants – and supports a connection between herbivory and the ant-specific lineage of *Bartonella* identified by Russell *et al* (2009) in *Dolichoderus* and *Cephalotes.*

The much lower normalized bacterial abundances we observed in almost all other arboreal ants suggests that they have evolved fundamentally different ecological and symbiotic strategies for life in the canopy. Some, like the abundant and ecologically dominant genera *Azteca* and *Crematogaster* (Wilson 1987), may simply be less herbivorous. These taxa rarely hosted any visible gut bacteria in our visual surveys, had median normalized 16S rRNA gene concentrations two orders of magnitude lower than those of the high-abundance taxa, and showed microbial taxonomic profiles that were highly varied across individuals and colonies. As with previous findings, they also had somewhat more omnivorous stable isotope profile – 1 to 3‰ higher ∂^15^N ratios than in *Camponotus, Cephalotes,* and *Dolichoderus* – suggesting that they complement their predominantly low-N liquid diets (Davidson *et al.* 2004) with moderate amounts of animal protein. Consistent with a strategy that pushes the boundaries of nitrogen availability, these taxa are reported to have among the lowest overall biomass nitrogen content and the highest behavioral preference for nitrogen-rich over carbohydrate-rich foods (Davidson 2005). Both of these genera typically have large, fast-growing colonies with presumably high overall demand for nitrogen.

More puzzling, perhaps, are the arboreal ants that harbored very low densities of bacteria, but still maintained depleted ∂^15^N ratios in the same range as *Camponotus* and *Dolichoderus.* Some of these may acquire their nitrogen from specialized associations with myrmecophytic plants. For example, *Neoponera luteola* is an obligate associate of *Cecropia pungara* (Yu & Davidson 1997), and the colony we measured had the most herbivorous isotope signature of any ant in our dataset (Fig. S6). The specialized food rewards provided by this species of *Cecropia* are especially nitrogen-rich for the genus (Folgarait & Davidson 1995), and may provide a major proportion of the ant’s overall nitrogen budget. However, most of the *Pseudomyrmex* species we surveyed were not specialized residents of ant-plants, yet still had very low bacterial abundances and depleted ∂^15^N ratios (the one obligate mutualist species we did survey, *P. triplaris,* was similar in both respects). Paradoxically, arboreal *Pseudomyrmex* have also been reported to have relatively limited behavioral preferences for nitrogen-rich foods compared to other arboreal ants (Davidson 2005) or to ground-nesting congeners (Dejean *et al.* 2014), suggesting that they have not evolved particularly strong behavioral imperatives for nitrogen acquisition. If the low densities we observed in the guts of arboreal *Pseudomyrmex* truly correspond to a limited role for bacteria in these ants’ nitrogen economy, how should these foraging patterns be interpreted – as indications of adequate supply, or of limited demand? The relatively small colony size of free-living *Pseudomyrmex* species may simply require less nitrogen than the high-biomass colonies of *Azteca* and *Crematogaster.* Alternatively, arboreal *Pseudomyrmex* might form microbial associations at other lifestages (e.g. in the larval gut), or rely on alternative nitrogenous food sources, as in recent reports of fungal cultivation and consumption in the genus (Blatrix *et al.* 2012).

The comparative paucity of high bacterial loads among ground-nesting ants further supports the hypothesis that the extremely dense bacterial associations of some arboreal ants are adaptations to life in the canopy. In stark contrast to our findings for arboreal ants, ground-nesting ants showed no significant correlation between ∂^15^N isotope ratios and bacterial abundance – in fact, for visual estimates of abundance, there was a marginally significant trend towards higher densities in more carnivorous ants. Consistent with this trend, the only genera outside of *Camponotus, Dolichoderus,* and *Cephalotes* where we definitively observed gut bacteria in FISH micrographs were in the exclusively carnivorous army ants (Fig. 2).

Could bacterial associations facilitate extreme carnivory on the forest floor analogously to how they appear to have facilitated extreme herbivory in the canopy? Sequence-based surveys of bacteria have revealed consistencies among army ant microbiota (Funaro *et al.* 2011; Anderson *et al.* 2012; Lukasik *et al.* 2016; Russell *et al. In Press)* that seem to contrast with the highly variable communities that have been recovered from more generalist arboreal (Sanders *et al.* 2014) and terrestrial species (Lee *et al.* 2008; Ishak *et al.* 2011). While 16S rRNA sequencing was unsuccessful for most *Labidus* and *Neivamyrmex* individuals in our dataset, it did succeed for a large proportion of colonies and specimens from these two genera (and other army ants) that were collected elsewhere in the Americas (Lukasik *et al.* 2016). Specimens from that study commonly hosted gut bacteria from two army ant-specific groups – identified there as an undescribed Firmicutes lineage and an undescribed Entomoplasmatales lineage – which were also found in *Eciton* individuals characterized in this study. Biological roles of gut bacteria in carnivorous ants are not known; but high mortality in ants restricted to protein-rich foods – irrespective of carbohyrate content – also suggests a potential role for bacteria in ameliorating deleterious effects of obligate carnivory (Dussutour & Simpson 2012). If such associations do result in carnivorous ants hosting higher overall quantities of bacteria compared to more omnivorous species, the physiological demands of the association would appear to be satisfied by cell densities that are still orders of magnitude lower than in the canonical canopy-dwelling herbivores. More targeted investigations, using techniques with finer sensitivity at very low abundances, will be required to resolve this question.

### Evidence for intracellular bacteria in multiple ant lineages

Nutritive intracellular endosymbionts are common in a great variety of insects (Moran *et al.* 2008), but, with the significant exception of *Blochmannia* endosymbionts in the speciose genus *Camponotus,* surprisingly absent among ants. After the initial description of intracellular bacteria in the camponotini and some species of *Formica* (Blochmann 1888) over one hundred years ago (see also Dasch 1975), only very recently, with the discovery of a gammaproteobacterial endosymbiont in the invasive ant *Cardiocondyla* (Klein *et al.* 2015), have similar associations been described in other ant lineages. We found microscopic evidence suggestive of gut-localized intracellular bacteria in two other arboreal ant lineages, suggesting that these associations may be considerably more widespread in ants than was previously thought.

Putative bacteriocytes in one of the two colonies of *Myrmelachista* we examined (colony JSC-108) appeared similar to those of *Camponotus,* and like in *Camponotus,* contained large, rod-shaped bacteria. *Myrmelachista* are specialized twig-nesters and frequent inhabitants of ant-plants, which form specialized structures to house and sometimes feed the ant inhabitants. Relatively little is known about the ecology of most species in the genus (Longino 2006), though the association between *M. schumanni* and the ant plant *Duroia hirsuta* results in dense, almost agricultural stands of the host plant due to pruning activity of the ants (Frederickson *et al.* 2005; Frederickson & Gordon 2007). A study of another plant associate, *M. flavocotea,* whose colonies nest in species of *Ocotea,* showed that workers of this species have a stable isotope signature much higher than that of their host plant, suggesting a substantial degree of carnivory (McNett *et al.* 2009). The two colonies in our dataset had sharply divergent ∂^15^N isotope ratios: colony JSC-137 (*M. schumanni*), which we recovered from the ant plant *Cordia nodosa,* was at about 9‰ similar to values reported from *M. flavocotea.* The colony in which we observed putative bacteriocytes in worker midguts (JSC-108) had a much more herbivorous signature; at 3‰, among the lowest values we recovered in our dataset. This colony also had a higher median normalized 16S rRNA gene abundance by almost an order of magnitude (Fig. S6), and two of the three individuals sequenced had community profiles dominated by sequences assigned to the genus *Sodalis,* which has been described as an intracellular associate of flies and beetles (Moran *et al.* 2008). Sequences similar to *Sodalis* have also been reported from *Tetraponera* and *Plagiolepis* (Stoll *et al.* 2007; Wernegreen *et al.* 2009). While these observations are anecdotal, the correlated variation in presence of putative bacteriocytes, inferred diet, and overall bacterial abundance make *Myrmelachista* an attractive candidate for further study.

The potentially intracellular bacteria we observed in *Dolichoderus* were even more striking. Individuals in this genus consistently harbored dense concentrations of bacteria in a blanket-like band around the proximate portion of the midgut (Fig. 2). These cells were quite large, with irregular, often heavily branched, morphologies. Branching morphologies have been reported in *Blochmannia* (Buchner 1965), and large, irregular phenotypes in other endosymbionts result from runaway gene loss associated with the bottlenecks of vertical transmission (McCutcheon & Moran 2011). Interestingly, previous microscopy-based investigations failed to detect bacteriocyte associates in a Brazilian species, *Dolichoderus attebaloides,* instead suggesting a dense packing of bacterial cells between host midgut cells (Caetano *et al.* 1990) – an intimate but still extracellular habitat that would also be consistent with our observations. While the data we present here cannot verify an intracellular location, the ultrastructural position of these cells (near the outer margin of midgut tissue, rather than interfacing with the gut lumen) and their derived cellular morphology both strongly suggest an intimate relationship with the host.

### Most ants have very few bacteria

We were surprised by how few bacteria we found in most ants. How unusual are these numbers? Direct numerical comparisons with organisms from other studies are challenging. Absolute bacterial abundances are only rarely reported in the literature (Engel & Moran 2013). When they are, the techniques used to derive them vary significantly, making direct comparison suspect. Furthermore, insects scale in body size across many orders of magnitude, so some normalization by host insect size is necessary. Given these caveats, the bacterial loads we measured in most ants were quite low compared to other insects. Normalized by roughly estimated adult body weight (rather than to DNA concentration, for comparison to values from the literature), gut bacterial densities in low-abundance ants were on the order of 10^5^ (*Ectatomma* and *Gigantiops*) to 10^6^ (*Azteca* and *Crematogaster*) bacteria per gram, substantially lower than the ~10^8^ estimated per gram in *Drosophila* (Ren *et al.* 2007; Engel & Moran 2013). By contrast, higher-abundance ants (*Cephalotes, Camponotus,* and *Dolichoderus*) had closer to 10^9^ bacteria per gram – similar to values that have been estimated for aphids (Mira & Moran 2002), honey bees (Martinson *et al.* 2012), and humans (Savage 1977).

The shape of the normalized bacterial abundance distribution within colonies of these low-abundance ant species hints at fundamental differences in the mechanisms underlying host/microbiome relationships among ant taxa (Fig. S13). High-abundance ant genera tended to have more normal distributions of normalized bacterial abundance, and more consistent taxonomic profiles within colonies, implying that the loss of microbial cells through excretion and death is balanced in these taxa by cell division of autochthonous lineages in the host. By contrast, distributions in low-abundance taxa were heavily right-skewed: while the median individual in low-abundance ant genera typically had very few detectable bacteria, we occasionally found individuals with much higher densities. These skewed distributions are reminiscent of similar patterns in *Drosophila*, which exhibit rapid decreases in bacterial abundance when starved or transitioned to sterile media, suggesting that the dynamics of the gut microbiome are weighted towards extinction (Broderick *et al.* 2014).

Understanding the significance of these infrequent, high-titer individuals will likely be important to understanding the nature of ‘typical’ ant-microbe interactions: do they represent dysbiotic individuals, in which host suppression of bacterial growth has failed? Do they reflect the recent ingestion of meals containing high concentrations of bacteria? Ants have evolved numerous ways of suppressing unwanted microbial growth inside their nests, including antibiotics derived endogenously from unique metapleural glands (Yek & Mueller 2010) and exogenously from specialized symbioses with actinomycete bacteria (Schoenian *et al.* 2011). This tendency towards microbial fastidiousness may extend to the inside of their guts, as well.

### On the importance of quantification in host-associated microbial ecology

Our findings highlight the utility of quantification methods as a complement to surveys of sequence diversity in host-associated microbiomes. Amplicon sequencing techniques describe relative, not absolute, differences in bacterial abundance. Consequently, comparisons may be easily made between samples with little or no awareness as to how they differ with respect to the total number of bacteria present – a variable that is likely to be profoundly relevant to biological interpretation. In our study, the associations we observed between bacterial abundance and major ecological variables of habitat and stable isotope composition (Fig. 5) were not apparent in a PCoA ordination of bacterial diversity (Fig. 4). In mammals, even convergently-evolved herbivores can host communities of largely similar microbes, leading generally to clear grouping of samples by host ecology (Ley *et al.* 2008; Muegge *et al.* 2011). In ants, the relationships between species with dense populations of gut bacteria and their symbionts appears to be largely idiosyncratic to the host genus, increasing the difficulty of identifying broad correlations from diversity data alone. Had our analysis been limited to 16S rRNA amplicon sequencing, we would have found much more limited evidence to support an association between gut bacteria and arboreal herbivory in rainforest ants.

**Fig. 5:**
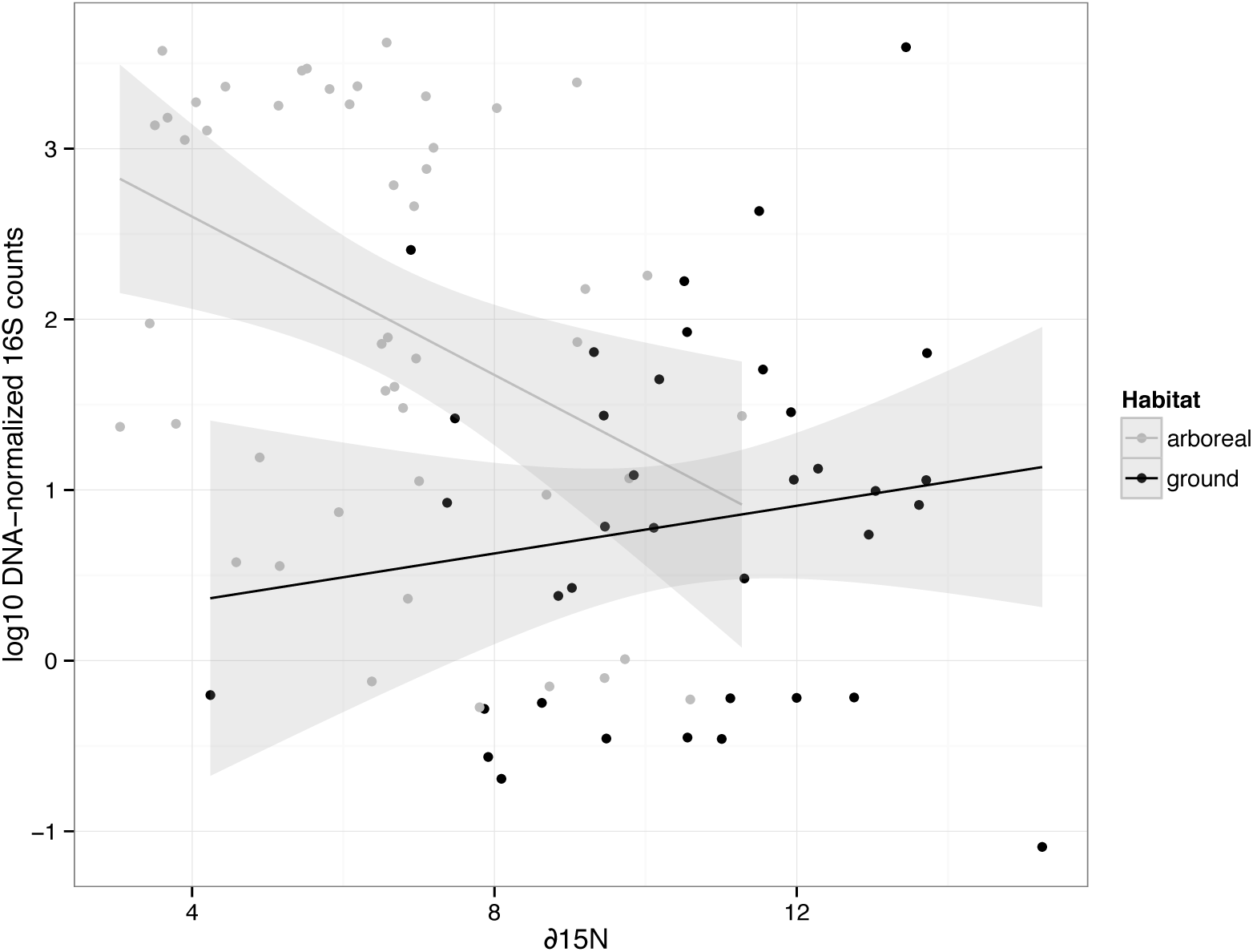
Normalized bacterial abundances (log_10_ qPCR 16S rRNA copy number per picogram DNA) by stable nitrogen isotope ratio. Each point represents the median value for a colony. Separate linear regressions (+/- 95% CI) fit to arboreal and ground-dwelling ants. Note that the simple linear fit is for illustration only; slope estimates for the mixed model used in analysis are presented in Fig. S12.

Characterization of bacterial abundance in samples should also help to interpret potential technical confounds, such as the presence of contaminant amplicons derived from reagents, the relative contribution of which should be inversely correlated to the original amount of bacteria in the sample (Salter *et al.* 2014). These challenges are likely to be especially relevant in small-bodied insects with variable bacterial populations, like many of the ant genera we observed in this study. Even without considering variance in host-normalized bacterial densities, ants from the same colony can span orders of magnitude in body size, leading to large differences in template quantity when amplifying from individuals. In practice, we have observed that within-colony variance community similarity is especially high in genera described here as having low overall bacterial abundances (Fig. S10) (Lee *et al.* 2008; Ishak *et al.* 2011; Sanders *et al.* 2014). Although the present study focuses on gross differences among ant genera with high- and low-density bacterial associations, detailed studies of bacterial communities in lower-abundance insect guts will need to take extensive measures to avoid technical confounds associated with low input biomass, as illustrated by a recent survey of the gut communities of Argentine ants (Hu *et al. In Press*). For these studies, pairing amplicon-based community profiling data with direct estimates of absolute bacterial abundance will help to provide important biological context to sequence diversity information.

### Conclusion

The explosion of 16S rRNA gene sequencing studies has justifiably led to an explosion of interest in animal microbiota (McFall-Ngai *et al.* 2013). Sequencing gives easy access to information about the composition of microbial communities, leading to extraordinary insights into the function and diversity of host-associated bacteria. Here, by demonstrating that gut bacterial densities help to explain the relationship between diet and habitat in rainforest ants, we have shown that simply surveying the abundance of these microbes can be useful as well, providing insights into ecology and potential function that would not be obtained by sequencing alone. Importantly, the techniques we used the assess abundance here – quantitative PCR and fluorescence microscopy – still only tell part of the biological story of these communities. The DNA detected by both methods does not necessarily correspond to metabolically active, or even viable, cells. To fully elucidate the functional and ecological consequences realized from these differences in potential will require more detailed investigations that can measure activity in addition to counting cells.

Still, the differences we observed among ants span several orders of magnitude, suggesting the potential for major differences in the roles of bacterial populations at each end of the spectrum. We humans host about a kilogram of bacteria in our gut (“Microbiology by numbers.” 2011); a *Cephalotes* ant, scaled to human size, would harbor roughly the same amount. The bacteria in the gut of *Gigantiops destructor,* similarly scaled, would weigh about as much as a roast coffee bean. We posit that these differences in magnitude correspond to differences in physiology with major relevance to the host.

## Acknowledgements

We thank Frank Azorsa and Stefan Cover for assistance with identification of specimens; Gabriel Miller, Lina Arcila Hernandez, Antonio Coral, and the staff of CICRA for assistance with collections and field work; Stephen Worthington and Kareem Carr for statistical advice; and Peter Girguis for providing laboratory facilities and support. Funding for this work was provided in part by a Putnam Expedition grant and NSF DDIG to JGS, and a Japanese Society for the Promotion of Science Short-Term Postdoctoral Fellowship, no. PE13061, to PL. MEF acknowledges the financial support of NSERC and the University of Toronto.

## Data Accessibility

Sequence data are available for analysis on the Earth Microbiome Project portal on Qiita, study number 10343 (https://qiita.ucsd.edu/study/description/10343), and deposited in the EMBL-EBI European Nucleotide Archive, accession number ERP014516. All other data and metadata, along with scripts necessary for generation of all data figures in this publication, are available in the Dryad Data Repository under accession [add accession here].

## Author Contributions

JGS and NEP conceived study; JGS, MEF, and PL performed field collections; JGS performed molecular work; NEP, RKo, JAR, and RKn contributed laboratory space and material; PL and RKo performed microscopy; JGS performed analyses; all authors contributed to writing manuscript.

## References

Anderson KE, Russell JA, Moreau CS et al. (2012) Highly similar microbial communities are shared among related and trophically similar ant species. Molecular Ecology, 21, 2282–2296.

Billen J, Buschinger A (2000) Morphology and ultrastructure of a specialized bacterial pouch in the digestive tract of *Tetraponera* ants (Formicidae, Pseudomyrmecinae). Arthropod Structure & Development, 29, 259–266.

Blatrix R, Djieto-Lordon C, Mondolot L et al. (2012) Plant-ants use symbiotic fungi as a food source: new insight into the nutritional ecology of ant-plant interactions. Proceedings Of The Royal Society B-Biological Sciences, 279, 3940–3947.

Blochmann F (1888) Ueber das regelmässige Vorkommen von bakterienähnlichen Gebilden in den Geweben und Eiern verschiedener Insecten. Zeitschrift fur Biologie, 24, 1–676.

Broderick NA, Buchon N, Lemaitre B (2014) Microbiota-induced changes in *Drosophila melanogaster* host gene expression and gut morphology. mBio, 5, e01117–14.

Buchner P (1965) Endosymbiosis of Animals with Plant Microorganisms (B Mueller, Tran,). John Wiley & Sons, Ltd.

Bution ML, Caetano FH, Zara FJ (2007) Comparative morphology of the ileum of three species of *Cephalotes* (Formicidae, Myrmicinae). Sociobiology, 50, 355–369.

Caetano FH, da Cruz-Landim C (1985) Presence of microorganisms in the alimentary canal of ants of the tribe Cephalotini (Myrmicinae): Location and relationship with intestinal structures. Naturalia, 10, 37–47.

Caetano FH, Tomotake M, Pimentel M, Mathias M (1990) Internal morphology of workers of *Dolichoderus attelaboides* (Fabricius, 1775)(Formicidae: Dolichoderinae). I. Digestive tract and associated excretory system. Naturalia, 15, 57–65.

Caporaso JG, Bittinger K, Bushman FD et al. (2010a) PyNAST: a flexible tool for aligning sequences to a template alignment. Bioinformatics, 26, 266–267.

Caporaso JG, Kuczynski J, Stombaugh J et al. (2010b) QIIME allows analysis of high-throughput community sequencing data. Nature Methods, 7, 335–336.

Caporaso JG, Lauber CL, Walters WA et al. (2011) Global patterns of 16S rRNA diversity at a depth of millions of sequences per sample. Proceedings Of The National Academy Of Sciences Of The United States Of America, 108, 4516–4522.

Caporaso JG, Lauber CL, Walters WA et al. (2012) Ultra-high-throughput microbial community analysis on the Illumina HiSeq and MiSeq platforms. The ISME Journal, 6, 1621–1624.

Cook S, Davidson D (2006) Nutritional and functional biology of exudate-feeding ants. Entomologia Experimentalis Et Applicata, 118, 1–10.

Dasch GA (1975) Morphological and molecular studies on intracellular bacterial symbiotes of insects. Yale University.

Davidson D, Patrell-Kim L (1996) Tropical arboreal ants: why so abundant? In: Neotropical Biodiversity and Conservation (ed Gibson AC), pp. 127–140. Neotropical Biodiversity and Conservation, Los Angeles.

Davidson DW (2005) Ecological stoichiometry of ants in a New World rain forest. Oecologia, 142, 221–231.

Davidson DW, Cook SC, Snelling RR, Chua TH (2003) Explaining the abundance of ants in lowland tropical rainforest canopies. Science, 300, 969–972.

Davidson D, Cook S, Snelling R (2004) Liquid-feeding performances of ants (Formicidae): ecological and evolutionary implications. Oecologia, 139, 255–266.

Dejean A, Labrière N, Touchard A, Petitclerc F, Roux O (2014) Nesting habits shape feeding preferences and predatory behavior in an ant genus. Naturwissenschaften, 101, 323–330.

Douglas A (2006) Phloem-sap feeding by animals: problems and solutions. Journal Of Experimental Botany, 57, 747–754.

Dussutour A, Simpson SJ (2012) Ant workers die young and colonies collapse when fed a high-protein diet. Proceedings Of The Royal Society B-Biological Sciences, 279, 2402–2408.

Edgar RC (2013) UPARSE: highly accurate OTU sequences from microbial amplicon reads. Nature Methods, 10, 996–998.

Eilmus S, Heil M (2009) Bacterial associates of arboreal ants and their putative functions in an obligate ant-plant mutualism. Applied and Environmental Microbiology, 75, 4324–4332.

Engel P, Moran NA (2013) The gut microbiota of insects - diversity in structure and function. FEMS Microbiology Reviews, 37, 699–735.

Engelbrektson A, Kunin V, Wrighton KC et al. (2010) Experimental factors affecting PCR-based estimates of microbial species richness and evenness. The ISME Journal, 4, 642–647.

Feldhaar H, Straka J, Krischke M et al. (2007) Nutritional upgrading for omnivorous carpenter ants by the endosymbiont *Blochmannia*. BMC Biology, 5, 48.

Folgarait PJ, Davidson DW (1995) Myrmecophytic Cecropia - Antiherbivore Defenses Under Different Nutrient Treatments. Oecologia, 104, 189–206.

Frederickson ME, Gordon DM (2007) The devil to pay: a cost of mutualism with *Myrmelachista schumanni* ants in “devil's gardens” is increased herbivory on *Duroia hirsuta* trees. Proceedings of the Royal Society of London B: Biological Sciences, 274, 1117–1123.

Frederickson ME, Greene MJ, Gordon DM (2005) Ecology: “Devil's gardens” bedevilled by ants. Nature, 437, 495–496.

Funaro CF, Kronauer DJC, Moreau CS et al. (2011) Army ants harbor a host-specific clade of Entomoplasmatales bacteria. Applied and Environmental Microbiology, 77, 346–350.

Hu Y, Holway DA, Lukasik P et al. By their own devices: invasive Argentine ants have shifted diet without clear aid from symbiotic microbes. Molecular Ecology, 1–56.

Hu Y, Lukasik P, Moreau CS, Russell JA (2014) Correlates of gut community composition across an ant species (*Cephalotes varians*) elucidate causes and consequences of symbiotic variability. Molecular Ecology, 23, 1284–1300.

Ishak HD, Plowes R, Sen R et al. (2011) Bacterial diversity in *Solenopsis invicta* and *Solenopsis geminata* ant colonies characterized by 16S amplicon 454 pyrosequencing. Microbial Ecology, 61, 821–831.

Jaffe K, Caetano F, Sanchez P et al. (2001) Sensitivity of ant (*Cephalotes*) colonies and individuals to antibiotics implies feeding symbiosis with gut microorganisms. Canadian Journal of Microbiology, 79, 1120–1124.

Klein A, Schrader L, Gil R et al. (2015) A novel intracellular mutualistic bacterium in the invasive ant Cardiocondyla obscurior. The ISME Journal.

Koga R, Tsuchida T, Fukatsu T (2009) Quenching autofluorescence of insect tissues for in situ detection of endosymbionts. Applied Entomology and Zoology, 44, 281–291.

Lee AH, Husseneder C, Hooper-Bùi L (2008) Culture-independent identification of gut bacteria in fourth-instar red imported fire ant, *Solenopsis invicta* Buren, larvae. Journal Of Invertebrate Pathology, 98, 20–33.

Ley RE, Lozupone CA, Hamady M, Knight R, Gordon JI (2008) Worlds within worlds: evolution of the vertebrate gut microbiota. Nature Reviews Microbiology, 6, 776–788.

Longino JT (2006) A taxonomic review of the genus Myrmelachista (Hymenoptera: Formicidae) in Costa Rica. Zootaxa, 1141, 1–54.

Lozupone C, Knight R (2005) UniFrac: a new phylogenetic method for comparing microbial communities. Applied and Environmental Microbiology,71, 8228–8235.

Lukasik P, Newton JA, Sanders JG et al. (2016) The structured diversity of specialized gut symbionts of the New World army ants. bioRxiv.

Martinson VG, Moy J, Moran NA (2012) Establishment of characteristic gut bacteria during development of the honeybee worker. Applied and Environmental Microbiology, 78.

McCutcheon JP, Moran NA (2011) Extreme genome reduction in symbiotic bacteria. Nature Reviews Microbiology, 10, 13–26.

McFall-Ngai M, Hadfield MG, Bosch TCG et al. (2013) Animals in a bacterial world, a new imperative for the life sciences. Proceedings of the National Academy of Sciences, 110, 3229–3236.

McNett K, Longino J, Barriga P et al. (2009) Stable isotope investigation of a cryptic ant-plant association: *Myrmelachista flavocotea* (Hymenoptera, Formicidae) and *Ocotea* spp. (Lauraceae). Insectes Sociaux, 57, 67–72.

Mira A, Moran NA (2002) Estimating population size and transmission bottlenecks in maternally transmitted endosymbiotic bacteria. Microbial Ecology, 44, 137–143.

Moran NA, McCutcheon JP, Nakabachi A (2008) Genomics and evolution of heritable bacterial symbionts. Annual Review of Genetics, 42, 165–190.

Muegge BD, Kuczynski J, Knights D et al. (2011) Diet drives convergence in gut microbiome functions across mammalian phylogeny and within humans. Science, 332, 970–974.

Price MN, Dehal PS, Arkin AP (2009) FastTree: computing large minimum evolution trees with profiles instead of a distance matrix. Molecular Biology and Evolution, 26, 1641–1650.

Ren C, Webster P, Finkel SE, Tower J (2007) Increased internal and external bacterial load during *Drosophila* aging without life-span trade-off. Cell metabolism, 6, 144–152.

Roche R, Wheeler D (1997) Morphological specializations of the digestive tract of *Zacryptocerus rohweri* (Hymenoptera: Formicidae). Journal Of Morphology, 234, 253–262.

Rognes T, Flouri T, Nichols B, Quince C, Mahé F (2016) VSEARCH: a versatile open source tool for metagenomics. PeerJ.

Rubin BER, Sanders JG, Hampton-Marcell J et al. (2014) DNA extraction protocols cause differences in 16S rRNA amplicon sequencing efficiency but not in community profile composition or structure. MicrobiologyOpen, 3, 910–921.

Russell JA, Moreau CS, Goldman-Huertas B et al. (2009) Bacterial gut symbionts are tightly linked with the evolution of herbivory in ants. PNAS, 106, 21236–21241.

Russell JA, Sanders JG, Moreau CS Hotspots for symbiosis: function, evolution, and specificity of ant-microbe associations from trunk to tips of the ant phylogeny (Hymenoptera: Formicidae). Myrmecological News, 1–27.

Salter SJ, Cox MJ, Turek EM et al. (2014) Reagent and laboratory contamination can critically impact sequence-based microbiome analyses. BMC Biology, 12, 87.

Sanders JG, Powell S, Kronauer DJC et al. (2014) Stability and phylogenetic correlation in gut microbiota: lessons from ants and apes. Molecular Ecology, 23, 1268–1283.

Savage DC (1977) Microbial ecology of the gastrointestinal tract. Annual Review Of Microbiology, 31, 107–133.

Schmitt-Wagner D, Friedrich MW, Wagner B, Brune A (2003) Phylogenetic diversity, abundance, and axial distribution of bacteria in the intestinal tract of two soil-feeding termites (*Cubitermes* spp.). Applied and Environmental Microbiology, 69, 6007–6017.

Schoenian I, Spiteller M, Ghaste M et al. (2011) Chemical basis of the synergism and antagonism in microbial communities in the nests of leaf-cutting ants. Proceedings of the National Academy of Sciences, 108, 1955–1960.

Stoll S, Gadau J, Gross R, Feldhaar H (2007) Bacterial microbiota associated with ants of the genus *Tetraponera*. Biological Journal Of The Linnean Society, 90, 399–412.

Thermo Scientific (2007). Detection of DNA with Quant-iTTM PicoGreen^®^ dsDNA Reagent in microplate format. 1–3.

Tobin JE (1991) A neotropical rainforest canopy, ant community: some ecological considerations. Ant-plant interactions, 536–538.

Walters W, Hyde ER, Berg-Lyons D et al. (2016) Improved Bacterial 16S rRNA Gene (V4 and V4-5) and Fungal Internal Transcribed Spacer Marker Gene Primers for Microbial Community Surveys. mSystems, 1.

Wilson EO (1987) The arboreal ant fauna of Peruvian Amazon forests: a first assessment. Biotropica, 19, 245–251.

Wolschin F, Holldobler B, Gross R, Zientz E (2004) Replication of the endosymbiotic bacterium *Blochmannia floridanus* is correlated with the developmental and reproductive stages of its ant host. Applied and Environmental Microbiology, 70, 4096–4102.

Yek SH, Mueller UG (2010) The metapleural gland of ants. Biological Reviews, 86, 774–791.

Yu DW, Davidson DW (1997) Experimental studies of species-specificity in *Cecropia-*ant relationships. Ecological Monographs, 67, 273–294.

Microbiology by numbers. (2011) Microbiology by numbers. Nature Reviews Microbiology, 9, 628.

